# Reactivating the complement system using peptidic bacterial labeling tag

**DOI:** 10.1101/2023.07.09.548267

**Authors:** Yael Belo, Einav Malach, Zvi Hayouka

## Abstract

The immune system plays a critical role in protecting the host against pathogens, including bacteria, viruses, and parasites. However, pathogens have evolved mechanisms to evade the immune system, for example by altering their surface proteins or by producing enzymes that can interfere with the immune response. These evasion strategies enable pathogens to escape detection and destruction by the immune system, which allows them to establish serious infections. Thus, there is a critical need for new strategies for developing antimicrobial agents. Here, we describe a novel strategy for targeting pathogens, by labeling them with a general peptide functioning as a bacterial binder, conjugated to a protein tag recognizable by the complement system, thereby activating the immune system against the target pathogen. To that end, we screened several pathogenic bacteria to find complement-resistant bacterial strain. A selected peptide binder was crosslinked with the C3b complement protein using glutaraldehyde. We show by an ELISA assay that the resulting complex binds the C5 complement protein with high affinity. We posited that by binding C5, this complex will be capable of initiating the alternative complement downstream proteolytic cascade, thereby inducing the formation of the membrane attack complex. Using this methodology, we were able to eradicate 90% of complement-resistant *E. coli* bacterial cells. By showing enhancement of complement sensitivity in complement-resistant pathogens, this work demonstrates the basis for new therapeutic approach capable of targeting pathogenic bacteria and activating the immune system against them.

## Introduction

Although the vast majority of bacterial strains are not harmful, there fewer than one hundred species that are pathogenic and capable of causing serious infectious diseases. The frequent use of wide-range antibiotics to treat these diseases contributes to the rise of antimicrobial resistance, which in turn leads to high hospitalization rates, significant patient suffering, illness, and death [1]. The protection of the host against invading pathogens relies heavily on the crucial and varied functions of the immune system [2], [3]. The human immune system has developed an elaborate network of cascades to fight microbial intruders [4], [5]. Within this network, the complement system acts as an important mediator in the elimination of pathogens, by constituting one of the earliest and most significant responses to pathogen infections [6], [7]. Complement activation occurs through one of its three activation pathways, namely, the classical, alternative and lectin pathways [8]. According to the alternative complement cascade, C3 protein is activated by C3 convertase, through cleavage of a single peptide bond of C3 into C3a and C3b protein fragments [9], [10]. As a result of the cleavage, C3b carries a thioester reactive group, which attaches to bacterial cell membranes by covalently binding hydroxyl groups found on the membrane [11]. Due to rapid hydrolysis of the reactive thioester group, most active C3b molecules (90%) do not succeed in attaching foreign matter [12]. In addition, C3b can bind a Bb fragment and become a C3 convertase, which continues to cleave C3 and induces high attachment of C3b to bacterial surfaces [13], [14], [15]. As a result, C3b complexes are produced and form high-affinity C5 convertase enzymes together with C3 convertases [16], [17], [18]. The resulting high density of C3b molecules on complement-opsonized foreign matter surfaces, forms high affinity attachment sites for C5 [11]. C5 convertases cleave C5 to C5a and C5b fragments [19], and C5b activates the alternative complement cascade by collaborating with C6-C9 complement proteins to create the Membrane Attack Complex (MAC) [20]. MAC then induces pores formation by injecting itself into the bacterial cell membrane, resulting in cell lysis [21]. Complement-dependent bacterial eradication is one of the fastest ways to kill an invading pathogen bacterium. While phagocytosis killing and subsequent intracellular eradication takes between 30 min to 1 h to occur, both the coating of bacteria with C3b molecules and the MAC-directed killing is reached within minutes [5]. The fact that pathogenic bacteria have evolved mechanisms to resist steps of the complement cascade, strongly supports the significant role the complement response plays in human defense against pathogens [22], [23], [4]. Resistance to complement elements, is also associated with the composition of the bacterial lipopolysaccharide (LPS). LPS is a vital component of the outer membrane in gram-negative bacteria, contributing to its structural integrity [24]. It consists of three distinct domains; the membrane-embedded lipid A, core oligosaccharides, and the O-antigen polysaccharides. LPS molecules having all three regions were defined as smooth LPS, while those lacking the O-antigen were termed rough LPS [25]. Virulence traits might be affected by the length and structure of the O-poly-saccharides, the most notable one being a resistance to complement and phagocytosis, or to killing by macrophages and neutrophils. There are also Gram-positive bacteria that exhibit resistance to complement treatment [26]. For instance, *Staphylococcus aureus* secretes small evasion molecules and expresses extracellular complement binding protein and extracellular fibrinogen-binding protein [27]. These proteins effectively bind to C3b and impede the activity of C3 and C5 convertases, thereby inhibiting the complement cascade [28]. These findings demonstrate the urgent need to develop strategies that will enhance the immune response towards immune resistant pathogens [29]. One example is a recently developed biological synthetic approach termed antibody recruiting molecules (ARMs), which has the ability to mediate immune clearance through small molecules capable of enhancing antibody attachment to bacteria, disease-relevant cells or viruses [29], [30], [31]. For example, Whitesides et al. utilized a bifunctional polyacrylamide (pA–V–F) presenting vancomycin and fluorescein groups [32], [33]. The vancomycin groups recognize a structural component of the bacterial cell wall peptides terminated in D–Ala–D–Ala, while the fluorescein groups facilitated visualization of this binding using fluorescence. The polymer-labeled bacteria were effectively recognized by anti-Fluor antibodies that triggered interactions with macrophages, leading to the promotion of polymer-labeled bacteria phagocytosis. In a related study conducted by Rice and colleagues, a strategy was proposed to enhance complement activation specifically targeting *Neisseria gonorrhoeae* and *Neisseria meningitidis* bacteria [34]. These bacteria express glycans that inhibit complement activation by binding complement inhibitor factor H (FH). To counteract this, Rice and colleagues developed a novel approach by fusing the bacterial binding domains of FH with the Fc region of IgG. This fusion protein, referred to as FH-FC, demonstrated the ability to mediate complement-dependent killing in vitro and exhibited promising efficacy in animal models of both gonorrhea and meningococcal bacteremia. In a conceptually related study, Rooijakkers and colleagues employed a click chemistry reaction to conjugate purified complement C3b onto *E. coli* bacteria [35]. They reacted the thioester of purified C3b with DBCO-PEG4-maleimide, resulting in C3b-DBCO, which they reacted with bacterial LPS decorated with Keto-deoxyoctulosonate-N3, leading to the creation of C3b-*E. coli* conjugates. Subjecting the conjugates to human neutrophils directly triggered phagocytosis and the subsequent destruction of the bacterial cells through the activation of Complement receptor 1. Here, we have employed a novel approach that distinguishes itself from previous methods [29-32]. Here, instead of focusing on a specific protein found on the surface of a particular pathogen or a particular structural component of bacterial cell walls, our approach targets microbial membranes using a broader targeting strategy. We have developed a random cationic peptide binder comprising of Lysine and Isoleucine amino acids, which interacts effectively with the hydrophobic and anionic properties of bacterial membranes. Unlike small and specific molecules such as ARMs, our peptide binder, termed 10-mer IK, is a mixture of 1028 different peptides. We anticipate that this diversity will impede the development of resistance, which is often observed with small and specific molecules. Our strategy does not involve the recruitment of antibodies for immune activation. Instead, a 10-mer IK peptide bacterial binder was conjugated to C3b, a crucial complement protein. This allowed us to tag bacteria with the 10-mer IK-C3b complex, aiming to induce enhanced complement activation towards these labeled bacteria.

## Results

### C3b Conjugation to a Peptide Binder

Our group has developed a new type of antimicrobial peptides termed random peptide mixture (RPM), comprising hydrophobic and cationic amino acids. We have previously shown that they exhibits efficient antimicrobial activity towards both Gram-negative and Gram-positive bacteria, including antibiotic-resistant bacteria [36], [37], [38], [39]. According to our findings, the chain length of an RPM significantly affects its antimicrobial activity. We observed lower antimicrobial activity for 10-mer RPMs compared to 20-mer length RPMs, which displayed strong and broad antimicrobial activity [37]. Therefore, we hypothesized that 10-mer RPMs might serve as beneficial general peptide bacterial binder. Since an important component of the innate immune system is the complement-dependent assembly of a proteins complex on the bacterial membrane, which induces bacterial death, we aimed to mimic this system by conjugating one of its main proteins, C3b, to a 10-mer IK RPM, which served as a bacterial peptide binder. The 10-mer IK RPM comprised a 1:1 molar combination of Isoleucine and Lysine. We hypothesized that the high positive charge of this 10-mer IK RPM will allow increased binding affinities to bacterial membranes. We have synthesized 10-mer IK labeled with 5(6)-carboxyfluorescein and observed great binding affinities to *E. coli R* and *S* bacteria (Figure S1 in Supplementary information). In addition, we examined the effect of 10-mer IK on the growth of E. *coli S* bacteria (Figure S2 in Supplementary information), which confirmed minimal growth-inhibiting effect. Taken together, these findings underscored 10-mer IK applicability as a general bacterial binder with great binding affinities. Next, C3b was conjugated to 10-mer IK using glutaraldehyde. Glutaraldehyde is an example of homo-bifunctional cross linker that usually couples through amino groups, such as free α-amino groups and lysine □-amino [40], [41]. As can be seen in Figure 1, following conjugation, the two subunits of C3b were coupled together, with a variable number of the 10-mer IK binder molecules conjugated to one C3b molecule. This observation implied that each complex contains multiple possible binding sites for the target bacteria. Therefore, these multivalent complexes were hypothesized to demonstrate increased binding affinity to the target bacterial pathogen. Next, we examined the 10-mer IK-C3b complex binding to *E. coli* bacterial cells. As can be seen in Figure 2, the 10-mer IK-C3b-fluorescein complex binds *E. coli S* bacteria in a concentration-dependent manner.

**Figure 1:**
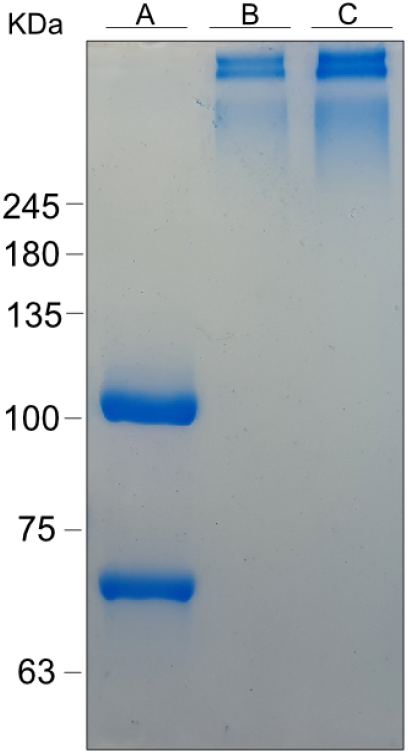
10-mer IK-Fluorescein labeled peptide conjugate to C3b protein. 10% SDS page of the conjugated 10-mer IK-C3b-Fluorescein. C3b molar mass is 176kDa. C3b composed of two chains, α chain is 101kDa and the β chain is 75kDa. A. C3b before conjugation to the IK peptide, runs as its two subunits. B. C3b conjugated to 10-mer IK-Fluorescein peptide. C. C3b protein conjugated to itself.

**Figure 2:**
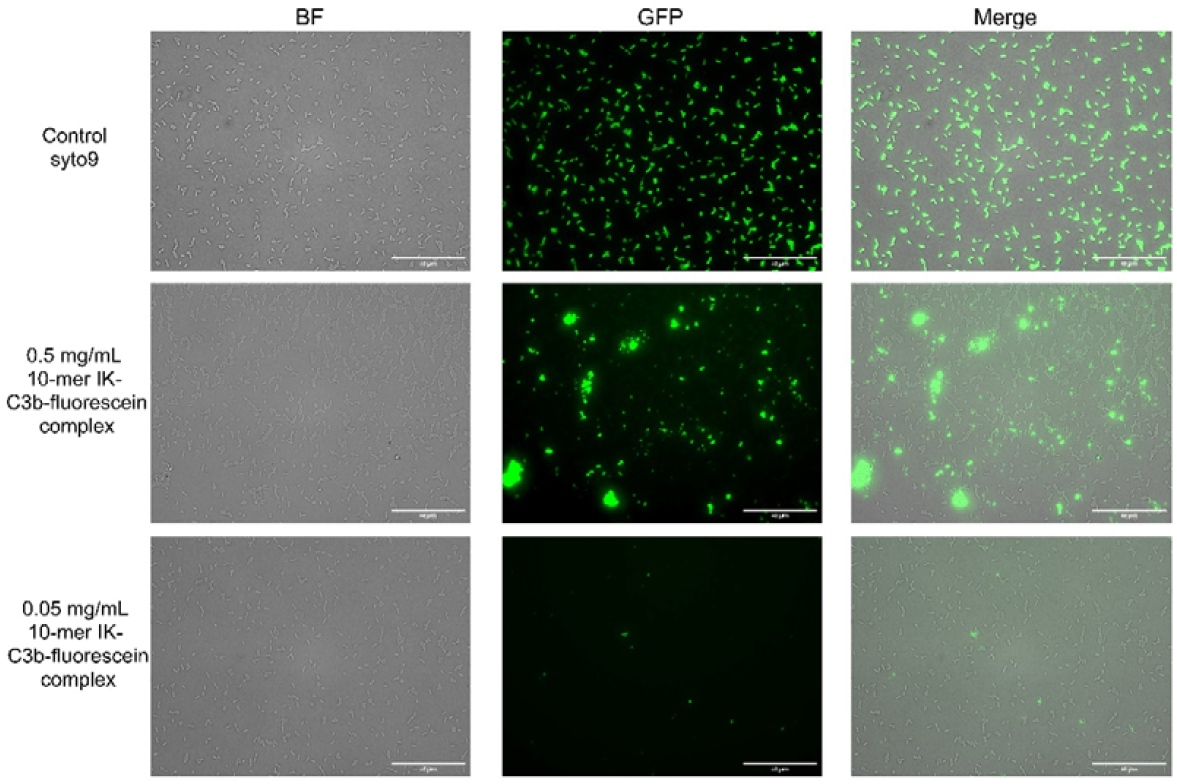
10-mer Fluorescein labeled IK-C3b binds *E. coli S*. EVOS microscope images of bound 10-mer Fluorescein labeled-IK-C3b complex to *P4-NR smooth E. coli* bacteria. Bacteria were grown to O.D. =1, and incubated together with the 10-mer IK-C3b-Fluorescein complex for 30 min at 37°C in PBS. A. Control sample which contained *P4-NR smooth E. coli* bacteria dyed with Syto9. B. Binding to 0.5 mg/mL 10-mer IK-C3b-Fluorescein complex. C. Binding to 0.05 mg/mL 10-mer IK-C3b-Fluorescein. Images shown were taken at a magnification of 40×.

### IK-C3b Complex Binds Complement C5

As noted, high density of C3b molecules on complement-opsonized foreign matter surfaces, forms high affinity attachment sites for C5 complement protein [11]. In addition, it was recently shown that at very high C3b density, MAC formation can be detected even in the absence of C5 activating enzymes [42]. To examine the capacity of the 10-mer IK-C3b complex to activate the complement alternative pathway, we conducted an enzyme-linked immunosorbent assay (ELISA). To that end, we coupled a biotin molecule to the 10-mer IK RPM and conjugated it to C3b protein using glutaraldehyde. C5 protein was coated onto ELISA microplate, and incubated with the 10-mer IK-biotin-C3b complex in serial dilutions. Binding level was assessed using streptavidin-HRP antibody. The 10-mer IK-C3b complex showed a specific and strong binding to C5 protein, with average K_*D*_ of 1.63 ± 0.53µM (Figure 3), in contrast to BSA or IK-IgG controls where no binding was observed. We concluded that 10-mer IK-C3b binds the C5 protein.

**Figure 3:**
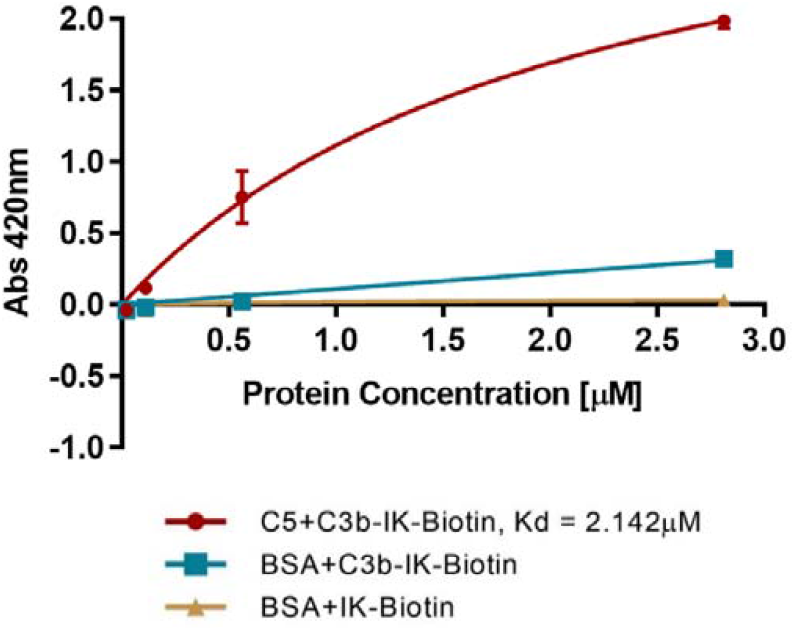
10-mer IK-C3b complex binds C5. C5 was fixated onto ELISA plate and C3b-IK-Biotin was added in a serial dilution (red). Mean ± SEM (n=2). Negative controls: blue: C5 was replaced with BSA, yellow: binding of IK-Biotin to BSA is shown (curve, overlaps with x-axis). Binding was examined using anti streptavidin-HRP antibody at 420nm. This is a representative result out of three biological repeats.

### Screening of Complement Resistant Pathogen Bacteria

To examine on a bacterial model the effects of innate immune system activation using a tagging strategy that employs our complex, we screened for bacterial strains that are either complement-resistant or complement-sensitive to act as controls. The *E. coli* strain *P4-NR smooth* (*E. coli S*), which possesses complement resistance, and the mutant *E. coli* strain *P4-NR*Δ *galU* (*E. coli R*), which is a rough strain and susceptible to the complement treatment

[25] were screened, as well as *PAO1, MRSA* and *E. coli K12*. We observed high resistant of *E. coli S, PAO1* and *MRSA* to serum treatment, and high sensitivity of *E. coli R* and *E. coli K12* (Figure 4).

**Figure 4:**
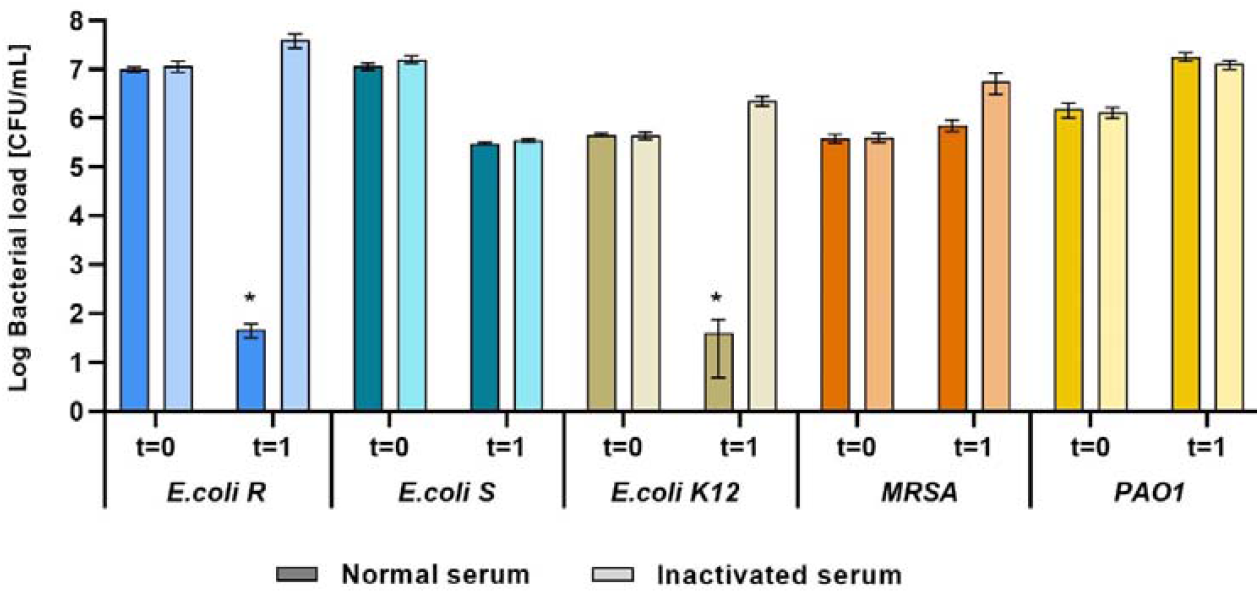
Screening of complement resistant pathogen bacteria. *E. coli P4-NR S, E. coli P4-NR R, E*.*coli K12, MRSA* and *PAO1* bacteria strains were incubated separately with normal bovine serum and inactivated serum for 1 hr at 37°C. Samples were micro diluted and plated on LB agar plates. Log of bacterial load at t=0, at the beginning of the experiment and after 1 hr of incubation. Mean ± SD (n=3). * P< 0.05, ** P< 0.01 (Unpaired t-test).

### IK-C3b Complex Induces Complement Sensitivity of *E. coli S* Bacteria

To evaluate the ability of the 10-mer IK-C3b complex in inducing clearance activity towards *E. coli S* serum-resistant bacteria, a serum sensitivity assay was conducted. In this assay, the rough strain was utilized as the positive control, while the smooth strain served as the negative control. Additionally, for bacterial tagging, the smooth strain was incubated with a 0.5 mg/mL concentration of the 10-mer IK-C3b complex. All bacteria samples were then incubated in 10% normal human serum or in heat inactivated serum as negative control. After 1 hr of incubation, microdilution was carried out, and each sample was plated on LB-agar plate. Finally, colonies count was performed to evaluate eradication effectiveness. The results showed complete eradication of the rough bacteria strain *E. coli R* (Positive control), whereas the smooth strain showed great resistance towards the serum (Figure 5A). In the presence of inactivated serum, no complement activity was observed. Incubating the smooth bacterial cells with the 10-mer IK C3b complex resulted in a higher eradication rate compared to the treatment with inactivated serum. The bacterial load (CFU/mL) of both tagged and untagged smooth bacteria treated with inactivated serum was quite similar, with the tagged bacteria at 100% and the untagged bacteria at 104.72% (Figure 5B). In a comparison between tagged *E. coli S* bacterial cells treated with inactivated serum, and those treated with normal serum, only 5.81% of the tagged cells survived the treatment. (Figure 5C). Following a 1 hr incubation period, the bacterial load of tagged *E. coli S* bacterial cells was found to be 10.51% of the untagged bacteria (Figure 5D). These findings demonstrated that the 10-mer IK-C3b tag is activating the complement system, in manner capable of enhancing the clearance of serum-resistant bacteria.

**Figure 5:**
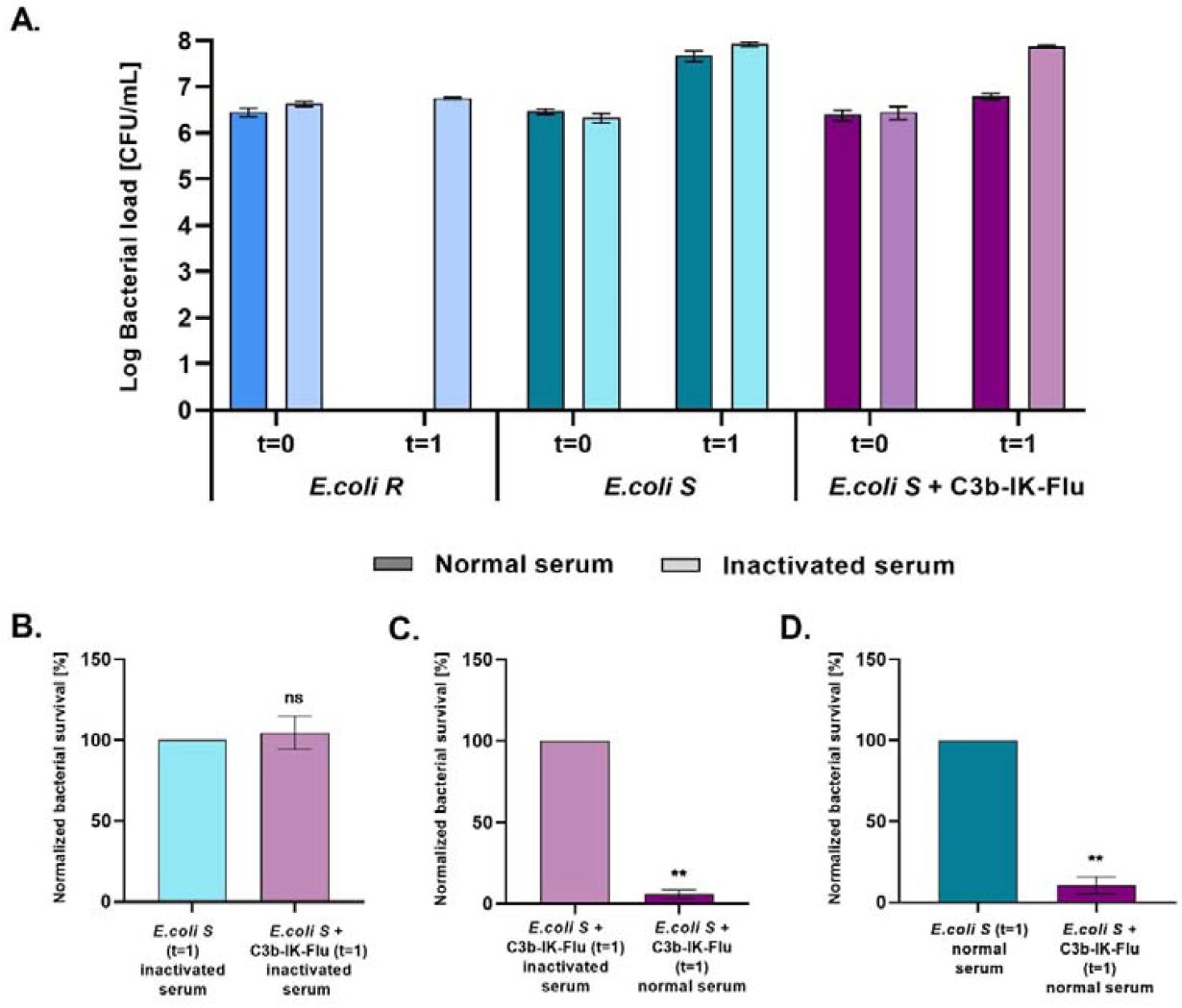
10-mer IK-C3b complex induces complement sensitivity of *E. coli S* bacteria. Bacteria were incubated with normal human serum (dark color) or inactivated serum (light color) for 1 hr at 37°C. Samples were micro diluted and plated on LB agar plates. A. Log of bacterial**S**l**u**o**p**a**p**d**le**o**m**f **e***E***n.ta***c***r***o***y***li* **d***R***a**,**ta***E. coli S* and *E. coli S* tagged with C3b-IK-Fluorescein, at t=0 and after 1hr of incubation in normal or inactivated serum (representative result from three biological repetitions). B. *E. coli S* and C3b-IK-Fluorescein tagged *E. coli S*, in inactivated serum. C. Tagged *E. coli S* in inactivated or normal serum. D. *E. coli S* and C3b-IK-Fluorescein tagged *E. coli S*, in normal serum. Mean ± SD (n=3). * P< 0.05, ** P< 0.01.

## Discussion

Our overarching goal was to test a versatile approach capable of tagging diverse types of cells, including bacteria and fungi, in order to activate the immune system against them. This ambitious goal was aimed at harnessing the power of the immune system to specifically target and combat a wide range of cell-based threats, irrespective of their nature or origin. In the current study we established a promising approach to redirect the immune complement response towards complement-resistant bacterial pathogens. The peptide binder we have selected, 10-mer IK RPM, was shown to possess strong binding affinity towards a diverse array of bacterial species, including both Gram-negative and Gram-positive bacteria. Consequently, this treatment demonstrates broad applicability across multiple bacterial species, rather than being limited to a single type. Furthermore, we recently revealed that RPMs exhibit a higher level of effectiveness in inhibiting the development of resistance, compared to antimicrobial peptides with specific sequence [43]. Here, by utilizing glutaraldehyde as a crosslinker, we observed a variable number of peptide binder molecules being conjugated to a single C3b protein molecule. We postulate that each such complex, therefore comprises multiple potential binding sites for the target bacteria. These multivalent complexes may thus have the potential to enhance the binding affinity for the targeted bacterial pathogen. We have further demonstrated that the 10-mer IK-C3b-fluorescein complex binds *E. coli S* in a concentration-dependent manner, and have provided evidence that the 10-mer IK-C3b complex exhibits good binding affinity to C5. Based on these observations, it is likely that our complex has activated the alternative pathway cascade, leading to the formation of the Membrane Attack Complex (MAC). We hypothesize that the abundantly-tagged bacteria with the IK-C3b complex, mimic a scenario in which there is a high density of C3b molecules present on complement-opsonized bacteria. This high density of C3b molecules usually creates an environment that facilitates the formation of high-affinity attachment sites for the C5 complement protein [11]. Finally, our findings demonstrate that *E. coli S* tagged with the 10-mer IK-C3b complex is significantly more susceptible to serum treatment, compared to untagged *E. coli S*. This suggests that the presence of the IK-C3b tag enhances the sensitivity of the bacteria to the bactericidal effects of the serum, possibly through increased complement-mediated killing. The new methodology presented, holds the potential to pave the way for novel treatments targeting diverse bacterial infections, prompting further research to explore and further utilize its application. Such ongoing research could contribute to the development of innovative therapies for combating bacterial infections and addressing the challenges posed by the ability of the bacteria to evade the immune system and to proliferate in the host.

## Materials and Methods

### Bacterial strains

*E. coli* strains *P4-NR*Δ *galU rough* (*E. coli R*) and *P4-NR smooth* (*E. coli S*) were kindly provided by Prof. Nahum Shpigel. *E. coli* strain *P4-NR*Δ *galU rough* (*E. coli R*) contains kanamycin resistance. All bacterial cells used in this study were stored in 25% glycerol at 80°C.

### Peptide Synthesis

Peptides were synthesized by a standard Fmoc-based solid-phase peptide synthesis (SPPS) on Rink Amide resin (substitution 0.6 mmol/gr), using a peptide synthesizer (Liberty Blue). Amino acids were dissolved with dimethylformamide (DMF) to a final concentration of 0.2 M. Fmoc deprotection step was conducted by 20% piperidine in DMF. At the end of the synthesis, the peptides were cleaved from the resin by adding a solution containing 95% trifluoroacetic acid (TFA), 2.5% double-distilled water (DDW) and 2.5% triisopropylsilane (TIPS) and stirred for 3 hours. The mixture was then filtered, and the peptides precipitated by the addition of 40 mL cold diethyl ether to the TFA solution and centrifuged. The supernatant was then removed and the peptide pellet dried, dissolved in 20% acetonitrile in DDW, frozen with liquid nitrogen and lyophilized. The synthesis was validated by MALDI-TOF mass spectrometry.

### C3b conjugation to 10-mer IK using glutaraldehyde

Human C3b protein was purchased from complement technology and mixed together with 10-mer random IK peptide mixture conjugated to a fluorescein molecule in molar ratio of 1:29.5. Glutaraldehyde was added over 1-2 min period, dropwise, and mixed gently with a magnetic stir for 1 hr at room temperature. After 1 hr 10 mM Glycine was added to stop the reaction. This was followed by overnight dialysis vs. 0.5 L PBS buffer at 4°C, froze in liquid nitrogen and kept in -80°C.

### 10-mer IK-C3b-Fluorescein binding to *E. coli* bacteria

*E. coli P4-NR (smooth)* bacteria were grown to O.D. = 1 and incubated with 0.5 mg/mL or 0.05 mg/mL C3b-10-mer IK-Fluorescein complex for 30 min at 37°C, 200 rpm. Three washes with PBS were performed. Each sample was placed on thin layer of 1% agarose which prepared on microscope slide. Syto9 was added to the control sample. Images were obtained using EVOS M5000 imaging system (Thermo Fisher Scientific).

### ELISA Assay

Biotin molecule was coupled to 10-mer IK and conjugation to human C3b protein (complement technology) using glutaraldehyde was performed as already described. Human C5 protein (complement technology) was dialyzed overnight against sodium bicarbonate buffer pH=9 and fixed on ELISA plate in concentration of 1 µM. Following that, it was incubated with 10-mer IK-biotin-C3b in serial dilution (highest concentration was 2.81 µM). The level of binding was assessed using α-streptavidin-HRP antibody in a dilution of 1:1000. TNB reagent was used with addition of 0.5 M H_2_SO_4_ and the absorbance at 420 nm was measured by plate reader (Tecan).

### Serum Sensitivity Test

Both bacteria strains, *E. coli* strains *P4-NR smooth (E. coli S)* and P4-NRΔ galU *(E. coli R)* were grown to O.D. = 1 (10^8^ CFU/mL) in LB media and washed three times with PBS. The smooth bacteria strain was incubated together with 0.5 mg/mL 10-mer IK-C3b complex for 30 min at 37°C, 200 rpm in PBS. After three washes with PBS, all samples were diluted to a concentration of 10^6^ CFU/mL. In a 24 wells plate 450 µL of bacteria in PBS were mixed together with 50 µL human serum. In every experiment, each sample had three wells which were dedicated for two types of serum – normal human serum (complement technology) and heat inactivated human serum for 30 min at 56°C. The plate was incubated for 1 hr at 37°C. At the end of the incubation, microdilutions were made for each sample in 96 wells plate, 10 µL drops were plated on LB agar plates and incubated overnight at 37°C. In the same way, microdilutions and plating were done also at t=0, before incubation.

## Figures

**Figure S1:**
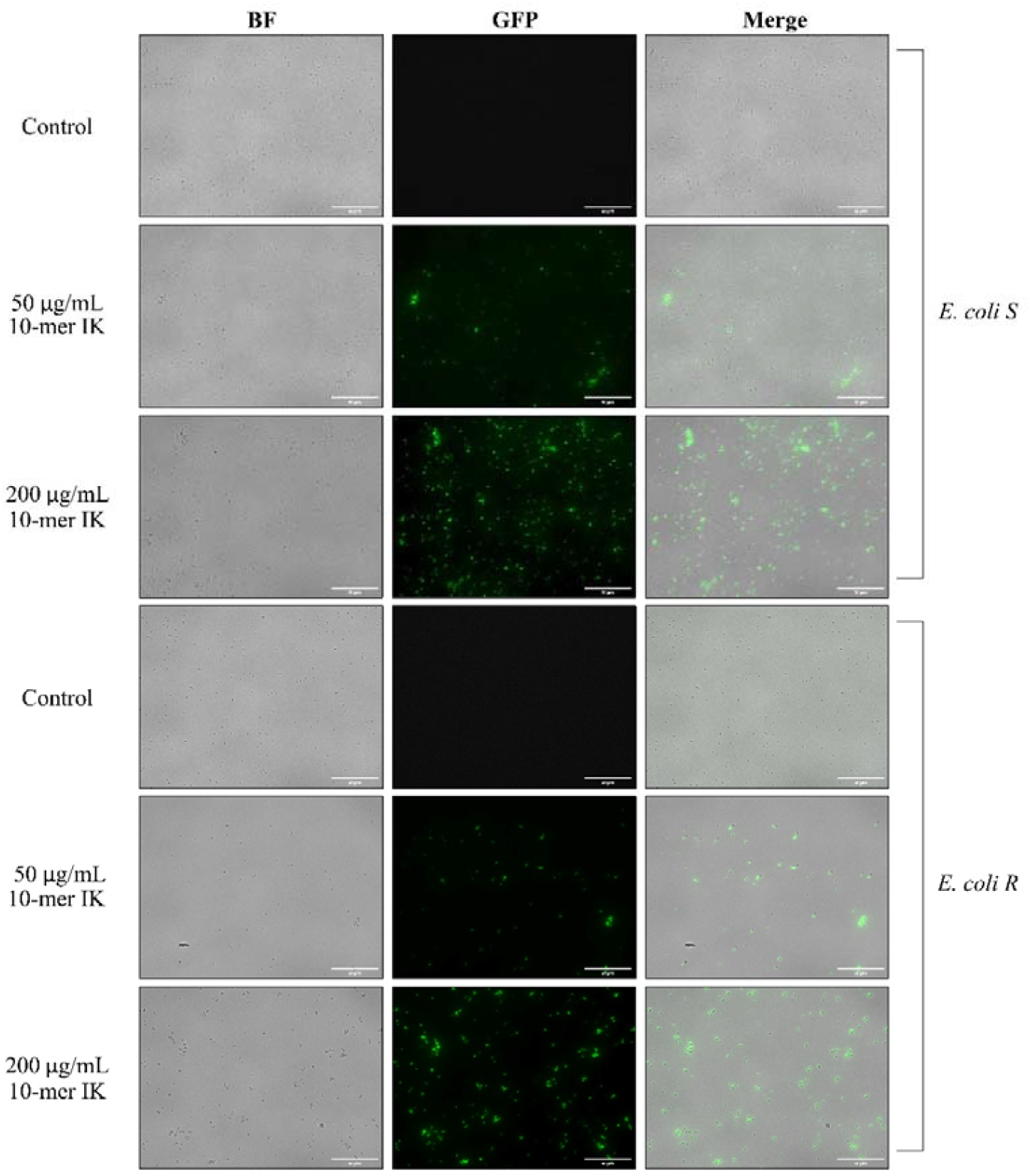
10-mer IK binds *E. coli* bacteria. EVOS microscope images of bound 10-mer IK *E. coli* S and *E. coli* R bacteria. Images shown were taken at a magnification of 40×. Bacterial cells were grown to O.D. = 0.1, and incubated with fluorescently labeled 10-mer IK for 30 min at 37°C in PBS.

**Figure S2:**
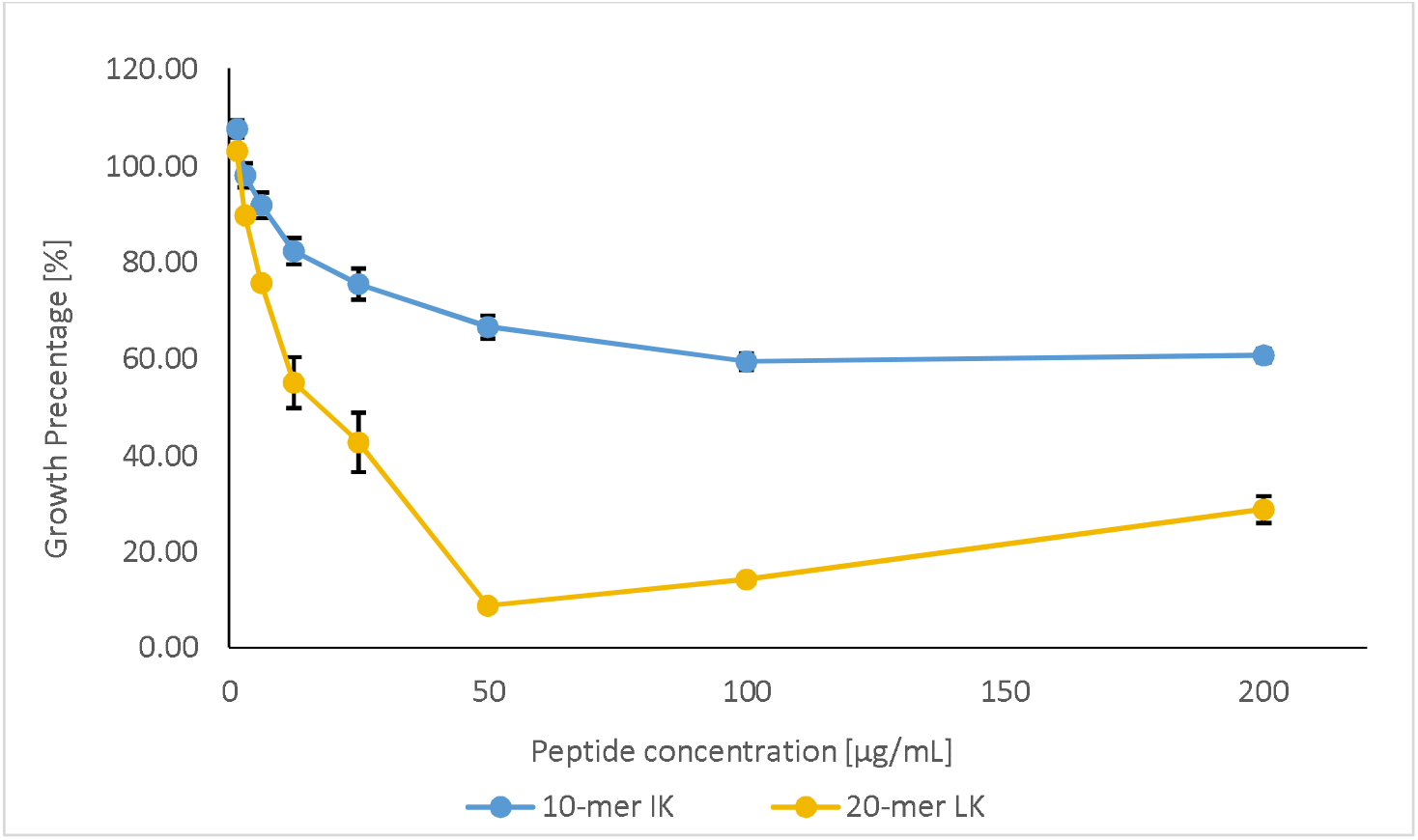
10-mer IK RPM showed low antimicrobial activity towards *E. coli*. Growth-inhibitory activities of 10-mer IK and 20-mer LK random peptide mixtures, toward P4-NR *E. coli* bacteria, after the incubation of 24hr at 37°C. The experiments were repeated three times (biological repeats) in triplicates (average ± SEM).

